# Calculating and Reporting Coefficients of Variation for DIA-based Proteomics

**DOI:** 10.1101/2024.09.11.612398

**Authors:** Alejandro J. Brenes

## Abstract

The Coefficient of Variation (CV) is a measure that is frequently used to assess data dispersion for mass spectrometry-based proteomics. In the current era of burgeoning technical developments, there has been an increased focus on using CVs to measure the quantitative accuracy of the new methods. Thus, it has also become important to define a set of guidelines on how to calculate and report the CVs.

This perspective shows the effects that the CV equation, as well as software parameters can have on data dispersion and CVs, highlighting the importance of reporting all these variables within the methods section. It also proposes a set of recommendations to calculate and report CVs for technical studies where the main objective is to benchmark technical developments with a focus on precision. To assist in this process a novel R package to calculate CVs (proteomicsCV) is also included.

## Introduction

Proteomics aims to characterise all proteins present in a specific cell, cell type, tissue or organism^1^. Recent breakthroughs have enabled a robust coverage of proteomes at scale, with setups detecting over 7,000 proteins in less than 30 minutes^2, 3^. Hence, the field is currently at an exciting stage where there is a drive to produce novel workflows and technological developments catered towards the high throughput analysis of low sample amounts, ranging to down to individual single cells. This has also produced a renewed interest in evaluating the quantitative precision of such workflows, instruments, or software packages, where a common practise is to measure and highlight the coefficient of variation (CV).

The CV is a measure of dispersion and a relative measure of variability. It compares the standard deviation to the measured mean. For proteomics in essence, it is comparing how close multiple measurements from different replicates are to each other. It is regularly used to assess the reproducibility of quantitative measurements, as the smaller the CV, the lower the deviation compared to the mean and the lower the dispersion, thus a lower CVs is equated to increased reproducibility and increased data quality. However a lower CV is not always positive, as it has been reported that inappropriate methodology such as faulty MS1 signal extraction or loose FDR can provide a lower but meaningless CV ^4^. As such it is important to avoid these pitfalls when using the CVs to evaluate the quantification.

Current developments focussed on technological advancements, including novel mass spectrometers^2, 5^, new sample handling and processing methods^5-11^ or updated software tools^12, 13^, frequently leverage CVs to evaluate the quantitative performance of their method. As such it would be beneficial to set up some guidelines on how to calculate and report the CVs for such studies. Using the Kawashima^14^ dataset, this perspective exemplifies how different the different CV formulas, data normalisation steps as well as software parameters affect the CVs produced. The reanalysis presented here are openly available at PRIDE^15^ under accession PXD052403. Using this same dataset a set of guidelines are suggested to calculate CVs in studies focusing on technical/technological developments and a different set of recommendations are made for biological studies. Furthermore, to assist in calculating the CVs in a standardised manner an R package (proteomicsCV) is provided to assist in the process.

### The effect of data normalisation and software parameters on CVs

Data normalisation is an important step in the proteomic data analysis pipeline, which has the objective to remove systematic bias, as such it also has a significant effect on the CVs that are produced and multiple tools have been developed to study this^16^. To visualise this effect, the Kawashima data was analysed in DIA-NN^17^ 1.8 setting the ‘Quantification strategy’ to high accuracy. The data was then analysed with and without a median normalisation strategy. As expected, the normalised data produces a median CV that is considerably lower than if CVs were calculated on the raw non-normalised intensity data (Fig. 1a). With this effect in mind, it is of clear importance to specify the normalisation procedures applied before calculating the CVs, and currently this is frequently omitted from the methods section. Furthermore, many software tools apply an array of data transformations by default, in some cases without users understanding that this is the case, which makes comparing CVs challenging.

**Figure 1.**
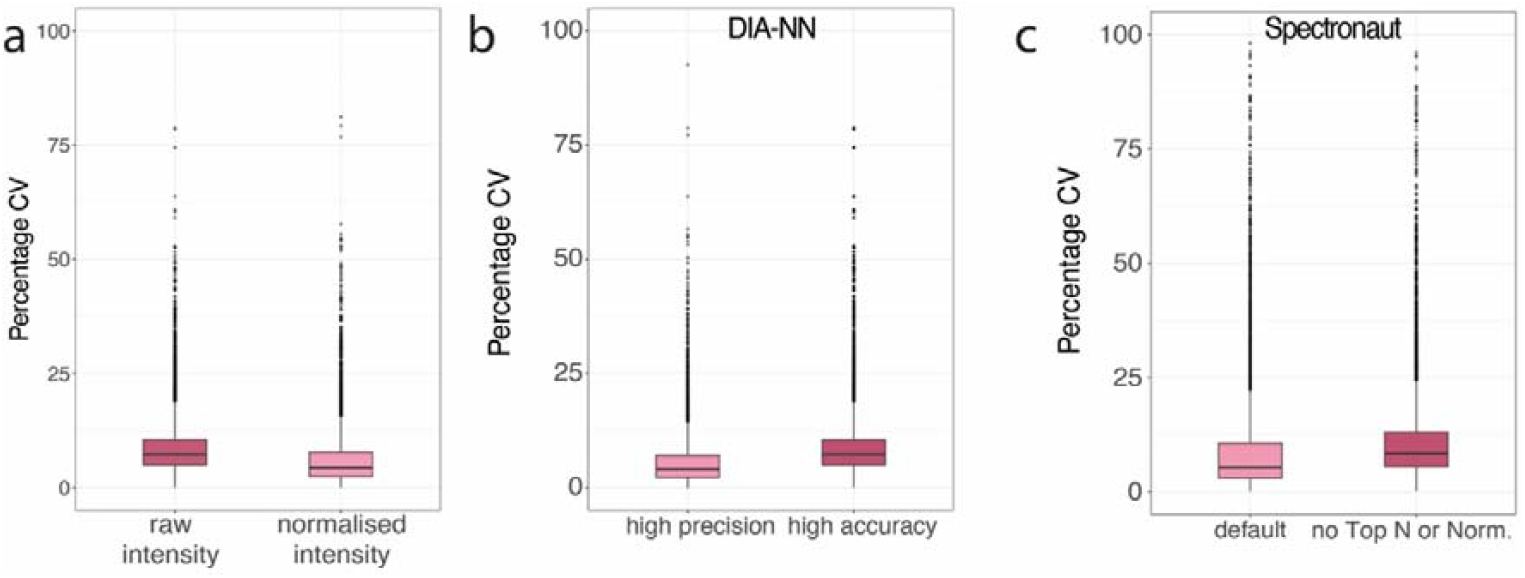
Normalisation effect on the CV and data distributions: **(a)** Boxplots comparing the Coefficient of Variation (CV) produced by using raw intensity compared to median normalised intensity. (**b)** Boxplots showing the CVs produced by DIA-NN when using the high precision vs high accuracy Quantification strategy. **(c)** Boxplots showing the CVs produced by Spectronaut when using the default vs stringent settings. The top whisker extends from the hinge to the largest value no further than 1.5 x IQR from the hinge; the bottom whisker extends from the hinge to the smallest value at most 1.5 x IQR of the hinge.

The automatic normalisation produced with DIA software is not tool specific and can occur both in DIA-NN^17^ and Spectronaut^18^. When processing DIA data with DIA-NN 1.8, there are two quantification options, ‘High Precision’ and ‘High Accuracy’. The ‘High precision’ option in the for quantification menu produces a median CV that is 45% lower than the high accuracy option (Fig. 1b). These results are due to a considerably less aggressive bias removal procedure that is used within the ‘High precision’ option. The results from the ‘High Precision’ option are very similar to the ‘High accuracy’ mode with an additional median normalisation step applied to the data (Fig. 1a). Understanding the differences between these two quantification modes in DIA-NN is an important consideration to comprehend and interpret the CVs that are produced. It should be noted new options are now available in DIA-NN 1.9 and the most similar alternative to ‘High Accuracy’ in DIA-NN 1.8 is the ‘legacy (direct)’ option.

Similarly, the default parameters that are pre-set in Spectronaut also include steps which minimise data dispersion and have an important effect on the CVs. The most important parameters which affect dispersion include a filter for the top 3 most abundant precursors and peptides, an operation that produces a CV almost as low as iBAQ^19^, as well as the default application of a global data normalisation step when the dataset is less than 500 raw files. This global normalisation is equivalent to the median normalisation step applied to the DIA-NN data (Fig. 1a). Disabling the Top 3 and data normalisation options from the default Spectronaut parameters produced a CV that is 35% higher (Fig. 1c), similar to the effect observed with DIA-NN with the two quantification modes. As with DIA-NN it is important to specify the parameter selection explicitly, specifying the Top N filter and normalisation instead of just stating the default parameters were used.

Thus, with the two most popular DIA software processing tools it is important to understand how the choice of parameters will affect the underlying data, and to report these specific parameters accordingly to help interpret the CVs that are produced. It is argued here that there are specific situations where using each of the parameter options can be beneficial, and others where it is not.

### Matching the CV formula to the appropriate data

Regardless of the type of study or normalisation step, it is important to understand how to calculate the CVs. There are two main formulas (see below) that are frequently used to calculate CVs for intensity based proteomic data. It is important to know when to it is appropriate to use each specific formula before applying it to the data.

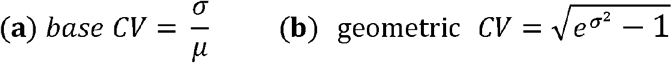

**CV Formulas: (a)** Base CV formula where σ is the standard deviation and µ is the mean protein intensity across the samples. **(b)** geometric CV formula where σ is the standard deviation of the log converted variance and *e* is Euler’s number (∼2.718).

Mass spectrometry-based intensity data are not normally distributed (Fig. 2a). The intensity only becomes approximately normally distributed after performing a logarithmic transformation (fig. 2b). Hence a common error is to apply a logarithmic conversion to the intensity data, while using the base CV formula on log transformed data. This erroneous use of the CV formula produces dramatically different results than when applying it to the non-log transformed intensity. The resulting median CV can be >14 times lower than that of the raw intensity with 75% of all proteins detected with a CV < 0.75% (Fig. 3a). This result is an artificial compression of the dispersion and does not faithfully represent the data acquired.

**Figure 2.**
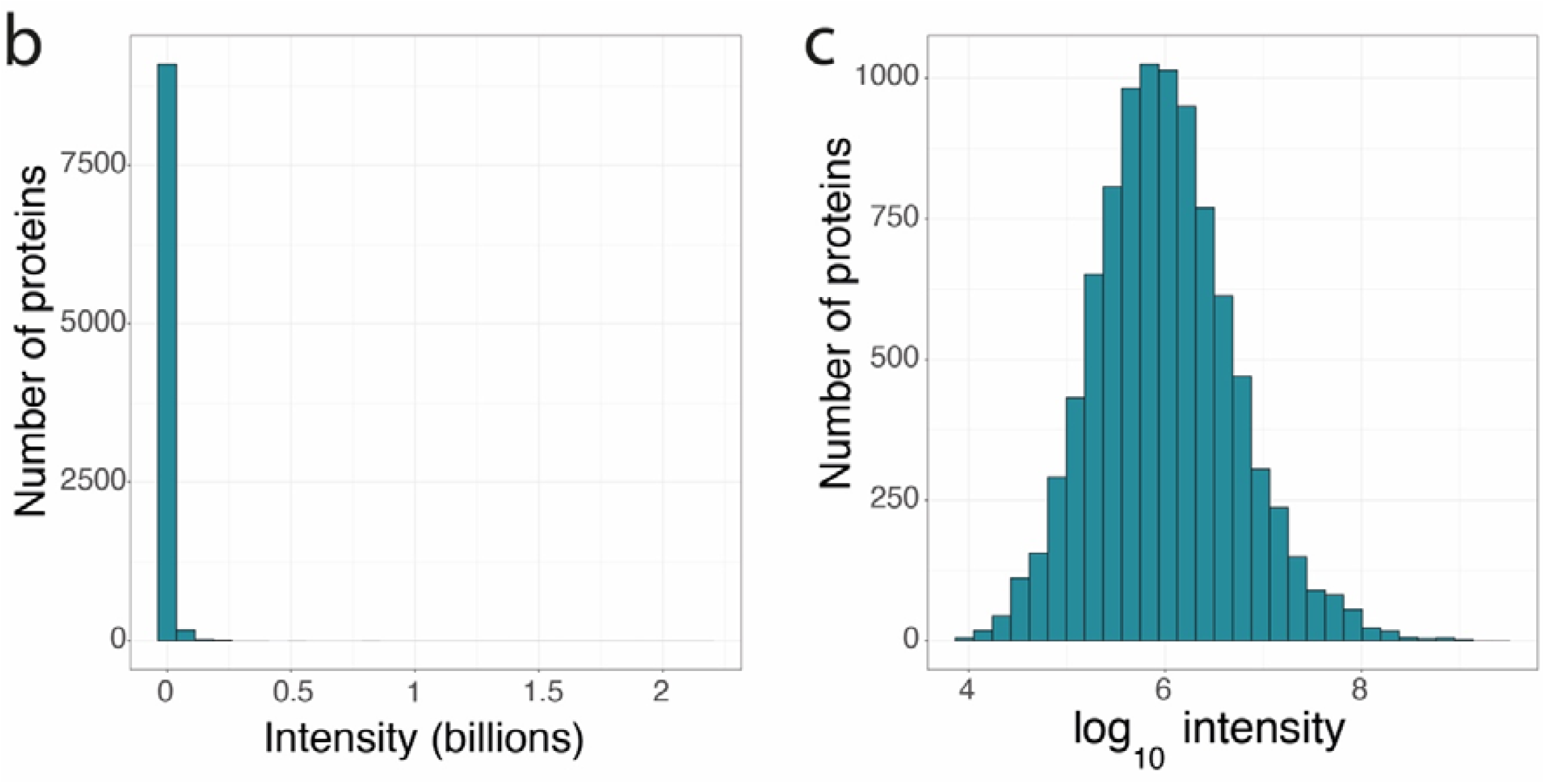
Log transforming intensity data: **(a)** Histogram showing the distribution of the raw intensity data. (b) Histogram showing the distribution of the log_10_ transformed intensity data.

**Figure 3.**
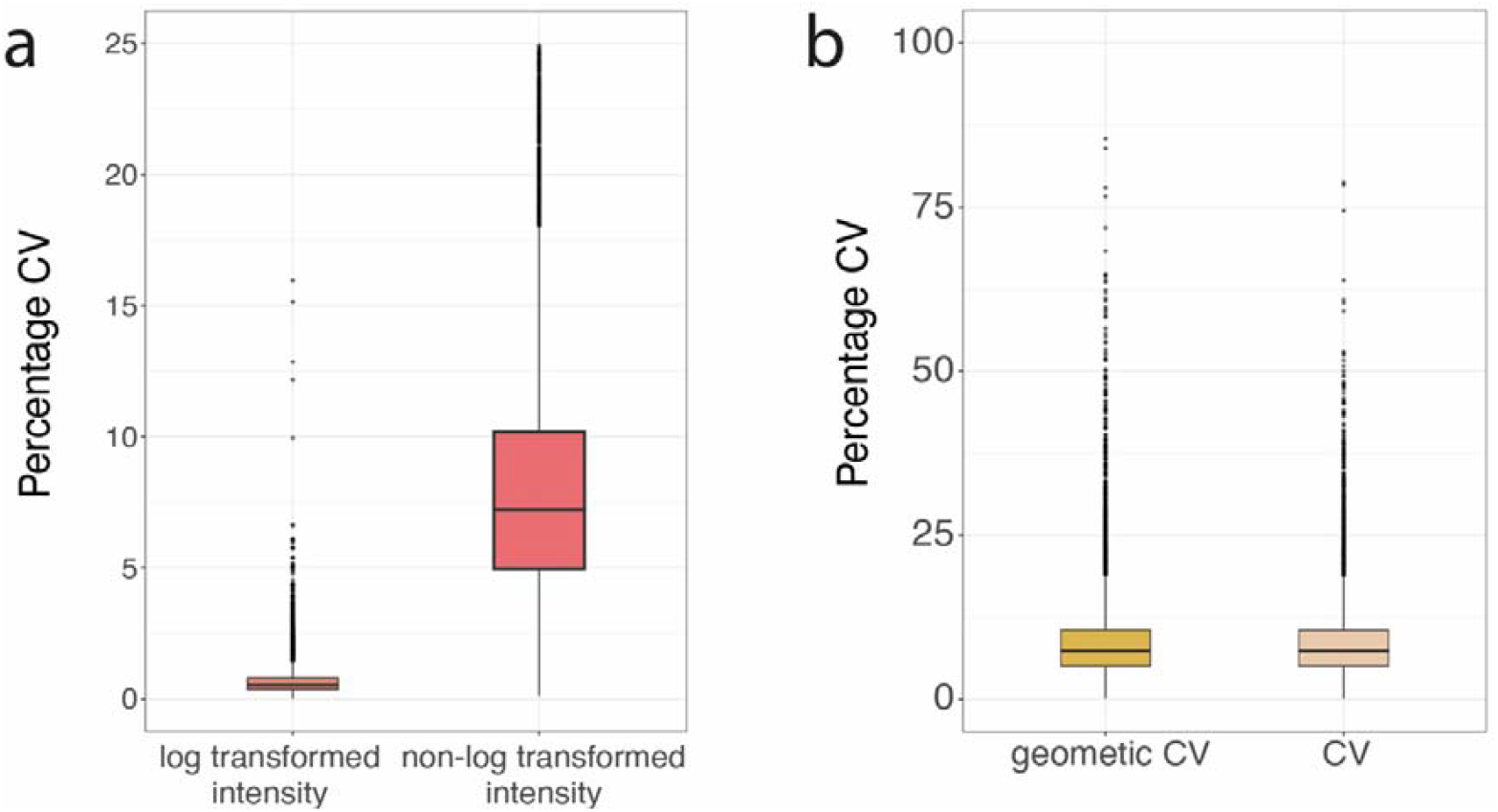
CVs across formulas: **(a)** Boxplots comparing the Coefficient of Variation (CV) produced by using the base formula with log transformed intensity and raw intensity. Y axis is cut at 25% to aid data visualisation. **(b)** Boxplot showing the CVs produced by using log transformed data calculated with the geometric CV formula and raw intensity data calculated with the base CV formula (CV). For all boxplots, the bottom and top hinges represent the 1st and 3rd quartiles. The top whisker extends from the hinge to the largest value no further than 1.5 x IQR from the hinge; the bottom whisker extends from the hinge to the smallest value at most 1.5 x IQR of the hinge.

To avoid this artificial shrinkage, using the base formula with log transformed data should be avoided. Hence for proteomic experiments the base CV formula should be applied to non-log transformed intensity. The geometric CV formula can also be utilized but on log transformed data. When both the previously formulas are correctly applied to the right type of data, the median CV produced are virtually identical and are more reflective of the underlying data (Fig. 3b).

### ProteomicsCV: an R package to calculate CVs

In an effort to simplify the CV calculation process for mass spectrometry-based proteomics data, an R package called “proteomicsCV” has been created and made openly available via the Comprehensive R Archive Network (CRAN). The package has a function to calculate the CVs for log transformed data, ‘protLogCV()’, and one function to calculate the CVs on non-log transformed data ‘protCV()’. The package can be easily installed by running the following command in R: “install.packages(‘proteomicsCV’)”.

### Summary and recommendations

This perspective focusses on the use of CVs for proteomics. Firstly, it is important to highlight that CVs themselves are a controversial measure, as they provide no insights into accuracy, nor the linearity of the quantitation and with inappropriate parameters can also produce low CVs that do not reflect high data quality, as such they should be used with caution and overreliance on them is not recommended^4^. To assess the quantitative performance and quality control of proteomics experiments multiple options exist beyond simply relying on CVs^20-22^.

However, it is undeniable that CVs are frequently used in proteomics. Thus, for studies which make use of CVs this perspective provides some tools and recommendations. Firstly, all studies should specify how the resulting CVs were calculated. Hence the methods section should state the formula that was used to calculate the CV as well as any data normalisation or data transformation steps that were performed before the CVs were produced. It is strongly recommended that the CVs are either calculated using the non-log transformed intensities with the base formula, or the log transformed intensities with the geometric CV formula.

The remaining recommendations differ for technical/technological studies and for biological experiments. For technical studies whose purpose is to benchmark the reproducibility in the quantification of a new instrument or method, and where the samples measured are mostly technical replicates it is recommended to use the non-normalised intensity data with no additional transformations or normalisation to calculate the CVs. This it will provide the most realistic overview of the dispersion present in the data.

Therefore, for these technical studies, specific parameters for DIA-NN and Spectronaut are recommended. When using DIA-NN <1.9 it is suggested to select the ‘high accuracy’ mode within the ‘Quantification strategy’ dropdown menu. For DIA-NN 1.9 onwards it is recommended to use the ‘legacy (direct)’ option. When using Spectronaut it is recommended to alter the default parameters by disabling the Top N (with 3 as default) for both major and minor groupings, and to disable the automatic data normalisation step. All the previously mentioned settings should be specified within the methods section of the relevant publications. The normalisation and transformations can minimise the differences in instrument performance, such as negating an increase in total intensity detected, therefore the raw intensity can be the most informative input when analysing performance in these technical studies.

For experiments aiming to explore the biology of different cells, cell types or conditions, with different biological replicates and whose primary objective is not to test reproducibility of the data produced by the mass spectrometer, then it is sensible to calculate the CVs on the normalised data. Biological replicates are expected to have increased variation, as protein expression can change considerably between individuals, compared to technical replicates which are expected to be more homogenous. Hence calculating the CVs on normalised data will provide a better overview of the data that will be used to analyse the biological dimension. As such it is reasonable to use ‘High Precision’ within DIA-NN 1.8. Furthermore, it is highly recommended to use the new QuantUMS options instead of the ‘legacy (direct)’ with DIA-NN 1.9. Similarly, it is also perfectly reasonable to enable normalisation within Spectronaut. However, the use of these parameters should be clearly stated within the methods, so readers are aware of the transformation applied before the CVs are calculated, and like with technical studies the formula used to calculate the CVs should be clearly stated in the methods.

This perspective provides some recommendations to improve the cross-comparability of CVs, especially for technical studies. Recent studies on similar instrumentation platforms display vastly different CVs, likely reflecting distinct data processing steps before they are calculated, however most of the studies do not document the process sufficiently to determine if this is the case. This perspective aims to assist researcher in avoiding this scenario. Furthermore, within this perspective an R package (proteomicsCV) is provided to assist in the process of correctly calculating the CVs for proteomics data by making it easier to understand when each CV formula should be used. By applying the correct formula and specifying the details of how the CVs were calculated, end users will be better informed to interpret the results. Furthermore, by providing software parameter recommendations for technical studies, it is hoped that this will make the results more comparable and representative.

## Data availability

The raw files, the FASTA file, the processed search results from DIA-NN using the two distinct quantification strategies and the two distinct Spectronaut searches are available on ProteomeXchange^23^ via PRIDE^15^ under the identifier PXD052403 (https://www.ebi.ac.uk/pride/archive/projects/PXD052403)

## Code availability

The R package proteomicsCV is freely available on CRAN https://cran.r-project.org/web/packages/proteomicsCV/index.html and can be installed in R using the ‘install.packages(“proteomicsCV”)’ command.

## Acknowledgments

The author would like to thank Vadim Demichev, Michael MacCoss, Alexey Chernobrovkin and Tanveer Batth for insightful twitter discussions about the nature of CVs and their use in proteomics.

